# Transcriptome analysis reveals the main metabolic pathway of c-GMP induced by salt stress in tomato seedlings

**DOI:** 10.1101/2020.10.28.358846

**Authors:** Xiaolin Zhu, Meifei Su, Xiaohong Wei, Yu Long, Baoqiang Wang, Xian Wang

## Abstract

Tomato is a model crop, as well as important food worldwide. In the arid areas, aggravation of soil salinity has become the primary problem that threatens the high yield in tomato production. As a second messenger substance, cyclic guanosine monophosphate (c-GMP) plays an indispensable role in plant response to salt stress through regulating cell processes to promote plant growth and development. However, this mechanism has not been fully explored in tomato seedlings. In this experiment, the tomato seeds were cultured in distilled water (CK), 20 μM c-GMP (T1), 50 mM NaCl (T2), 20 μM c-GMP + 50 mM NaCl (T3). The results show that 20 μM c-GMP effectively alleviated the inhibition of 50 mM NaCl on tomato growth and development, inducing the expression of 1580 DEGs. 95 DEGs were up-regulated and 442 DEGs were down-regulated (CK vs T1), whereas in the T2 vs T3 comparison 271 DEGs were up-regulated and 772 DEGs were down-regulated. Based on KEGG analysis, the majority of DEGs were involved in metabolism; exogenous c-GMP induced significant enrichment of pathways associated with carbohydrates, phenylpropanoids and fatty acid metabolism. Most *PMEs, acCoA, PAL, PODs, FADs*, and *AD* were up-regulated, and *GAPDHs, PL, PG, BXL4*, and *β-G* were down-regulated, which reduced susceptibility of tomato seedlings to salt and promoted their salt adaptation. The application of c-GMP promoted soluble sugar, flavonoids and lignin content, reduced accumulation of MDA, and enhanced the activity of POD. Thus, our results provide insights into the molecular mechanisms associated with salt tolerance of tomato seedlings.

## Introduction

Soil salinization is one of the most serious abiotic stresses that affects crop growth and development and yield production [1–3]. About 30 percent of agricultural land in the world is affected by high salinity [4]. The adverse effects of salinity on plants mainly include ionic toxicity, osmotic disturbance and oxidative stress, which may damage the integrity of cell membrane, affect cell metabolism, inhibit photosynthesis, and produce reactive oxygen species (ROS) [5–6].

In the natural environment, salt stress is mainly induced by high concentrations of Na^+^ and Cl^-^ in the soil. Excessive salt ions increase the osmotic potential of soil solution, causing dehydration of root cells, and affecting their physiological functions [7]. In addition, excessive concentration of salt ions inhibits uptake of other inorganic ions by roots, resulting in changes of the ion balance in plants. Ion imbalance and osmotic stress caused by salinity can inhibit the normal growth of plants, which in turn would affect crop yield and may even lead to death[8]. Therefore, how to alleviate salt stress and improve plant salt tolerance are critical in food production and in overcoming salinization.

Tomato (*Solanum lycopersicum*) is an annual species in family Solanaceae. Cherry tomato, known as ‘saint fruit’, is a popular type with good flavor. It is important food and has great economic value [9]. Applying exogenous substances to promote salt tolerance of plants is a relatively frequently-applied practice. Previous study [10] investigated the effects of exogenous Ca^2+^ on tomato germination and seedling growth under NaCl stress, indicating that exogenous substances can enhance the salt tolerance of tomato by influencing physiological and biochemical processes and antioxidant enzyme activities. The paper [11] found that exogenous ABA inhibited tomato seed germination by reducing the malondialdehyde (MDA) level to overcome drought stress. In addition, Pan[12]studied the phenotypic, physiological, biochemical, and molecular parameters, exploring the mechanism of action of the *SlbZIP38* transcription factor in tomato salt tolerance and proposed a theoretical basis of the adaptation of tomato to salt stress. In recent years, functional genomics has flourished. Tomato has become typical solanaceous crop in studying salt tolerance in plants [13–14]. However, the molecular mechanisms of exogenous c-GMP on the stress resistance of tomato plants and the use of transcriptome sequencing technology to characterize salt tolerance of tomato seedlings have not been reported so far.

Cyclic guanosine monophosphate (c-GMP) is one of the cyclic nucleotide signaling systems, and it is an important secondary signaling substance in animals and plants [15]. It recognizes intercellular and intracellular signals and amplifies them, regulates physiological and biochemical reactions in plants [16], and also creates signal transduction pathways for various environmental factors, abiotic stresses, plant hormones and other signaling molecules. In addition, c-GMP can regulate the adaptation of plants to biotic and abiotic stresses. Studies showed c-GMP can mediate ABA-induced stomatal closure [17]. The c-GMP signal has NO-independent and NO-dependent regulation modes [15, 18] It has also been reported that c-GMP can induce the synthesis of H_2_O_2_, and that the feedback regulation may be involved in the above signal transduction pathways [19]. It was reported that c-GMP is essential for the formation of anthocyanins and flavonoids, and several genes encoding secondary metabolite flavonoid biosynthesis enzymes including chalcone synthase (CHS) in soybeans are induced by c-GMP [20]. c-GMP is regulated by protein kinase PKGs, and can regulate phosphodiesterases, as well as cyclic nucleotide-gated ion channels (CNGCs)[21]. In addition, it was found that in pepper (*Capsicum annuum*), c-GMP can adjust uptake of Na^+^ and K^+^ to maintain ion balance [22].

The c-GMP membrane-permeable analogs are divided into 8-Br-c-GMP, 8-nitro-c-GMP and 8-mercapto-c-GMP [23–25], which all regulate plant growth and development, for example, seed germination and seedling growth. As a signal substance, c-GMP can enhance the tolerance of plants to stress. In less than 5 s, salt stress and osmotic stress can rapidly induce an increase in endogenous c-GMP, which can be prevented by the GC inhibitor LY83583. The recent studies found that 8-br-cGMP can enhance salt tolerance of *Arabidopsis thaliana* by inducing the expression of glucose-6-phosphate 1-dehydrogenase [19, 26, 73]pointed out that under stress, the expression of genes related to plant growth and development induced by 8-Br-c-GMP changed significantly, which in turn, affected the primary, secondary and energy metabolism in plants. The application of 20 μM 8-Br-c-GMP significantly promoted salt tolerance during seed germination [19]. These findings suggest that c-GMP is involved in a variety of complex physiological and biochemical processes.

Transcriptome sequencing (RNA-seq) technology, as a means of studying gene expression, has been used widely in many species and for various topics. It has been used in studying salt tolerance genes and microRNAs as well as the responses of plants to abiotic stress [74], particularly salt stress [27–28]. For example, Wang [29] used RNA-seq technology to reveal the potential molecular mechanisms of *Camellia sinensis* in response to salt stress and identified many candidate genes for further studies. Wan [30]applied 200 μ M c-GMP in dryland cotton (*Gossypium aridum*) and used transcriptome sequencing analysis to determine that the early stage of high salt stress was related to the transcriptional regulation of signal transduction and abiotic stress-inducing genes, whereas in the later stages of salt stress, it was mainly through adversity adaptation and defense. Xu[31] found that the molecular mechanism of *Arabidopsis thaliana* in response to salt stress was induced differential expression of transcription factors related to photosynthesis, auxin and osmoprotectants (such as trehalose and proline), enhancing salt tolerance of seedlings. In addition, large-scale evaluation of transcriptome resources in plants such as *Hippophae rhamnoides subsp. Sinensis* and *Piper nigrum* L. was projected to be used in the generation of new varieties [32].

The RNA-seq technology provides a good platform for understanding the mechanisms of c-GMP regulation of tomato seedling growth under salt stress. However, the transcription analysis of tomato seedling under salt stress induced by c-GMP has not been reported. The aim of this study was to explore the molecular mechanisms of c-GMP related to carbon, phenylpropanoids and fatty acid metabolism under salt stress, and to provide a theoretical basis for understanding salt tolerance of tomato seedlings, and ultimately improve tomato production in the arid and semiarid regions.

## Materials and methods

Cherry tomato seeds (with no mechanical damage) were surface sterilized with 2% v/v NaClO for 3-4 minutes, rinsed 5-6 times with sterile water and imbibed in distilled water (CK), 20 μM c-GMP (T1), 50 mM NaCl (T2), and 20 μM c-GMP+50 mM NaCl (T3) for 48 h. The choice of concentrations and treatment duration has been verified in our previous experiments (Table 1). Subsequently, seeds were placed in a glass dish (φ=9 cm) containing two layers of filter paper, with 4 mL of CK, T1, T2 or T3 solution added and replaced every day. The culture was carried out at 26°C and 12 h light/12 h dark. The whole seedlings were sampled at 3, 5, 7, and 9 d for the determination of physiological indices and growth. The 7-day-old seedling leaves were used for transcriptome sequencing.

**Table 1.** Summary of transcriptome sequencing data of exogenously administered c-GMP tomato leaves under salt stress

| Sample | Clean Reads | Clean Bases | Total reads | Mapped reads | GC content | %≥Q30 |
| --- | --- | --- | --- | --- | --- | --- |
| CK-1 | 21713544 | 6487237402 | 43427088 | 40221267(92.62%) | 43.05% | 93.40% |
| CK-2 | 21623659 | 6452921062 | 43247318 | 39977137(92.44%) | 43.06% | 94.29% |
| CK-3 | 22794060 | 6815569062 | 45588120 | 42115620(92.38%) | 42.89% | 93.84% |
| T1-1 | 21029112 | 6269326286 | 42058224 | 38802343(92.26%) | 42.94% | 93.80% |
| T1-2 | 21181097 | 6329645388 | 42362194 | 38661596(91.26%) | 42.91% | 93.98% |
| T1-3 | 21758314 | 6496985242 | 43516628 | 40198075(92.37%) | 42.96% | 94.02% |
| T2-1 | 25920637 | 7739107858 | 51841274 | 47904815(92.41%) | 42.67% | 94.31% |
| T2-2 | 24553877 | 7330762280 | 49107754 | 45543301(92.74%) | 42.77% | 94.00% |
| T2-3 | 26085941 | 7790260320 | 52171882 | 48352404(92.68%) | 42.74% | 93.62% |
| T3-1 | 23737433 | 7089477572 | 47474866 | 43814847(92.29%) | 42.57% | 91.42% |
| T3-2 | 25190167 | 7539092862 | 50380334 | 44349697(88.03%) | 43.54% | 91.15% |
| T3-3 | 23496662 | 7010732682 | 46993324 | 43273712(92.08%) | 42.83% | 90.91% |

### Tissue Sample and RNA Isolation

From the fifty plants per treatment, three biological replicates were established by randomly grouping three sets of plants in each treatment. Leaves were sampled 7 days after treatment commenced (CK, T1, T2, and T3), yielding 12 samples in total (1 time point and organ ×4 treatments ×3 biological replicates). The 12 samples were frozen immediately in liquid nitrogen and stored at −80°C before RNA isolation. Total RNA was isolated using a Trizol kit (Invitrogen, Carlsbad, CA, USA) and purified through a Qiagen RNeasy Column (Qiagen, Hilden, Germany) according to the manufacturer’s instructions. 1% agarose gel was run to check the integrity of RNA, The RNA integrity numbers (RIN) for the samples analyzed were 7.87, 8.03, 7.73, and 8.1 (leaves).

### Transcriptome sequencing and analysis of the results

After removing adaptors, the unknown nucleotides and those of low quality were filtered from the raw reads to obtain clean reads, which were then mapped onto the cherry tomato reference genome. Gene expression levels were calculated using FPKM values (fragments per kilobase of exons per million fragments mapped) by the cufflinks software 63. Genes with a false discovery rate (FDR) <0.005 and |log2 (FC)|≥1 were defined as differentially expressed genes (DEGs). The DEGs were subjected to KEGG pathway enrichment analyses using KOBAS 2.0. Furthermore, the mapman system was used to classify the transcripts at 0 and 20 μ M c-GMP.

### Determination of physiological parameters

Soluble sugar content was determined by anthrone colorimetry [33]. The starch content was measured by the Gibon method [34]. Malondialdehyde (MDA) was measured using an MDA assay kit (Jiancheng Bioengineering Institute, Nanjing). The activity of peroxidase (POD) was assayed as described by Hameed [35]. The lignin content was measured by the bromoacetyl method [36], and the flavonoid content was determined by spectrophotometry [37]. The fresh weight was determined using 30 seedlings randomly selected from each of the three biological replicates on day 7. Fresh samples were placed in an oven at 105 °C for curing, and were then oven-dried at 80°C for 48 h until the constant weight.

### Quantitative RT-PCR analysis

Nine DEGs (Differentially expressed genes) were selected for validation using q-RT-PCR. Tomato ACT was an internal reference gene. The primers used in this study were shown in Supplementary Table S1.

First-strand cDNA synthesis was performed using a PrimeScript TM RT reagent kit with gDNA Eraser (perfect Real Time), which removed genomic DNA. q-RT-PCR was performed with a Mx3000 P system (Applied Biosystems) using SYBR^®^ Premix Ex Taq™ II (TliRNaseH Plus) (TaKaRa) according to the manufacturer’s instructions. PCR amplification was performed in a 96-well plate according the following program: one cycle at 94 °C for 2 min, followed by 45 cycles at 94 °C for 5 s and 60 °C for 15 s. Melting curve analysis was implemented after incubation at 72°C for 10 s. Each PCR analysis was repeated at least three times. Relative gene expression amount was calculated using the threshold cycle 2^-ΔΔCt^ method [38].

### Statistical analysis

Parameters were statistically tested by analyses of variance, and comparisons of means were performed with the Duncan test (*P* < 0.05). Figures were prepared using the GraphPad Prism 5.0 software.

## Result

### 2.1 Exogenous c-GMP promoted tomato growth and enhanced salt tolerance

To confirm that c-GMP mitigated salt stress, tomato growth was monitored in the presence or absence of 50 mM NaCl after application of c-GMP. After 7 days of salt stress, the fresh and dry leaf biomass was higher in tomato seedlings treated with than without c-GMP (Figure 1A). Exogenous application of 20 μM c-GMP promoted the establishment and growth of cherry tomato seedlings. Compared with the control, root length, hypocotyl length, fresh weight, and dry weight of cherry tomato seedlings increased by 14.18%, 10.59%, 16.39%, and 9.72%, respectively, after treatment with c-GMP alone (T1) and decreased by 93.75%, 22.96%, 41.35%, and 33.85%, respectively, after treatment with NaCl (T2). Compared with T2, root length, hypocotyl length, fresh weight, and dry weight of cherry tomato increased by 94.29%, 34.59%, 35.08%, and 32.81%, respectively, after treatment with NaCl + c-GMP (T3). The effects of c-GMP on root length, hypocotyl length, fresh weight, and dry weight of cherry tomato under salt stress (T2-vs-T3) were significant. The germination of tomato seedlings was shown in Figure 1B.

**Figure 1.**
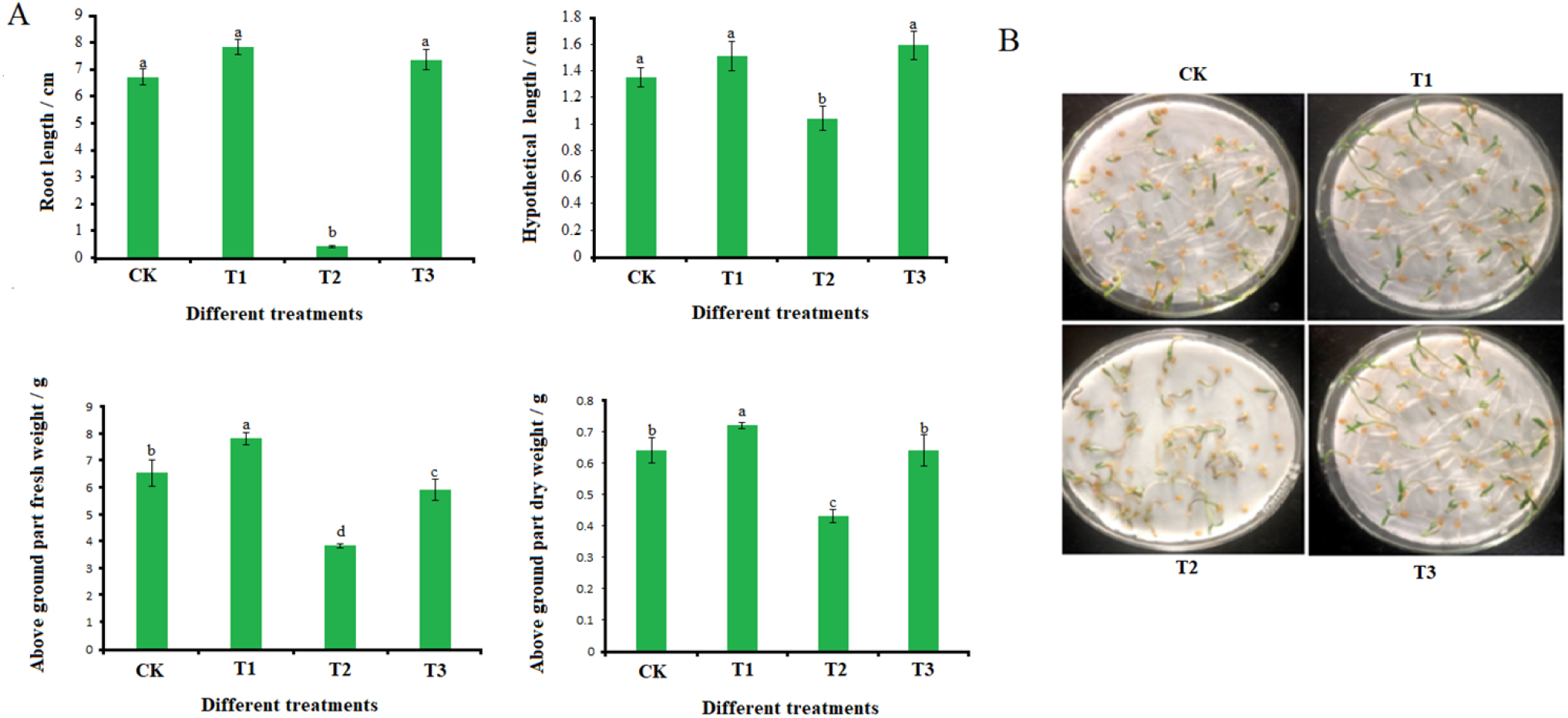
Plant growth, morphological, and physiological indicators. (A) Root length, hypocotyl length, fresh weight and dry weight of cherry tomato on the seventh day under different treatments; (B) Growth status of the seventh day under different treatments.

The error bar represents the standard error of the mean. The significant difference between different treatments was indicated by an sterisk when P⩽0.05.

Error bars denote standard error of the mean. Significant differences between varieties at p ≤ 0.05 are denoted by an asterisk.

### 2.2 Transcriptome sequencing and analysis of 12 samples

To identify genes associated with c-GMP-related alleviation of salt stress in cherry tomato seedlings, the Illumina high-throughput sequencing platform was used to sequence the cDNA libraries. A total of 83.35 G of clean data were obtained. The Q30 base percentage of each sample was not less than 90.91%. After mapping the clean reads obtained from the 12 libraries onto the tomato genome, the efficiency of the alignment varied from 88.03% to 92.74%, with multiple mapped reads found mainly in the rRNA or intergenic regions. Based on the alignment results, variable splicing prediction analysis, and gene structure optimization analysis, 1174 new genes were discovered, and 802 of them were functionally annotated (Table S2).

### 2.3 Identification of differentially expressed genes (DEGs)

To investigate the differentially expressed genes under control, salt stress and salt stress + exogenous c-GMP, we assessed all five comparisons (CK-vs-T1, CK-vs-T2, CK-vs-T3, T1-vs-T2, and T2-vs-T3). There were 537 and 1884 DEGs enriched in the 20 μM c-GMP and 50 mM NaCl treatments, respectively. Among them, 95 DEGs were up-regulated and 442 were down-regulated in the 20 μM c-GMP treatment, whereas 674 DEGs were up-regulated and 1210 were down-regulated in the 50 mM NaCl treatment (Figure 2B). A comprehensive list of differential genes is provided in Table 3S. Similar ratio changes in the up-regulated and down-regulated genes were observed under salt stress without (CK-vs-T2) and with exogenous c-GMP (T2-vs-T3).

**Figure 2.**
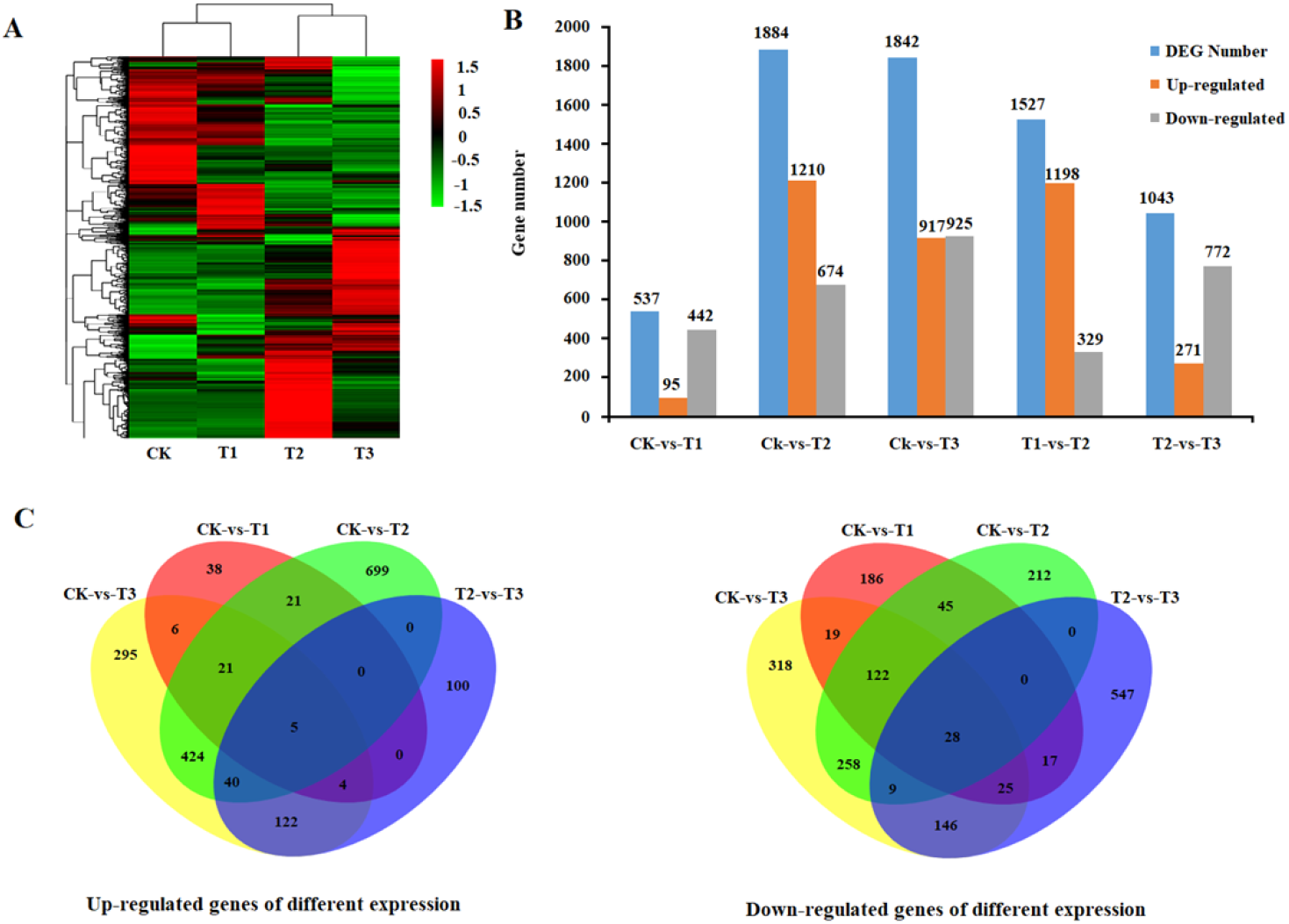
Identification of DEGs under four differential treatments. (A)The log2-fold-change value of the comparison process and the control, hierarchical clustering of DEGs was performed for different treatments to obtain four clusters; (B) Number of differential genes in comparison of four treatment combinations. (C) The venn plot shows the overlap between the differential genes in salt treatment, c-GMP treatment, and the combination of the two treatments.

The Venn diagram (Figure 2C, Table S4) showed there were 79 common differentially expressed genes between CK-vs-T1 and T2-vs-T3, including 9 up-regulated genes and 70 down-regulated genes. Total of 438 genes were specifically expressed under c-GMP treatment (CK-vs-T1). However, in the salt treatments (T2-vs-T3), the c-GMP-influenced DEGs increased to 944. It can be inferred that under salt stress, c-GMP induced expression of more genes to protect tomato seedlings from damage caused by unfavorable external factors.

### 2.4 Functional classifications of DEGs regulated by exogenous c-GMP under salt stress

All the annotated genes were mapped to the Kyoto encyclopedia of genes and genomes (KEGG) database. The KEGG pathway analysis provides classification of the complex biological functions of the genes [39]. Comprehensive analysis of differentially expressed gene in the CK-vs-T1 and T2-vs-T3 comparisons was provided in Table S5.

Top 20 KEGG enrichment analyses were shown in Figure 3. Adding 20 μM c-GMP caused 10 and 17 terms to be significantly enriched under 0 mM and 50 mM NaCl, respectively (Table S6). Several pathways, such as phosphatidylinositol signaling system, galactose metabolism and circadian rhythm - plants showed significant changes in CK-vs-T1, whereas phenylalanine metabolism, biosynthesis of terpenoids, diarylheptanes and gingerol, flavonoid biosynthesis, starch and sucrose metabolism, carbon metabolism, glycolytic/gluconeogenesis, amino acid biosynthesis, amino sugar and nucleotide sugar metabolism, fatty acid metabolism, steroid skeleton biosynthesis, pentose sugar and glucuronic acid conversion, glutathione metabolism, and steroid biosynthesis showed significant changes in T2-vs-T3 (corrected *P*<0.05). Analysis of the significantly enriched KEGG pathway caused by c-GMP under salt stress indicated involvements in the metabolism of carbohydrate, phenylpropanoids and fatty acids.

**Figure 3.**
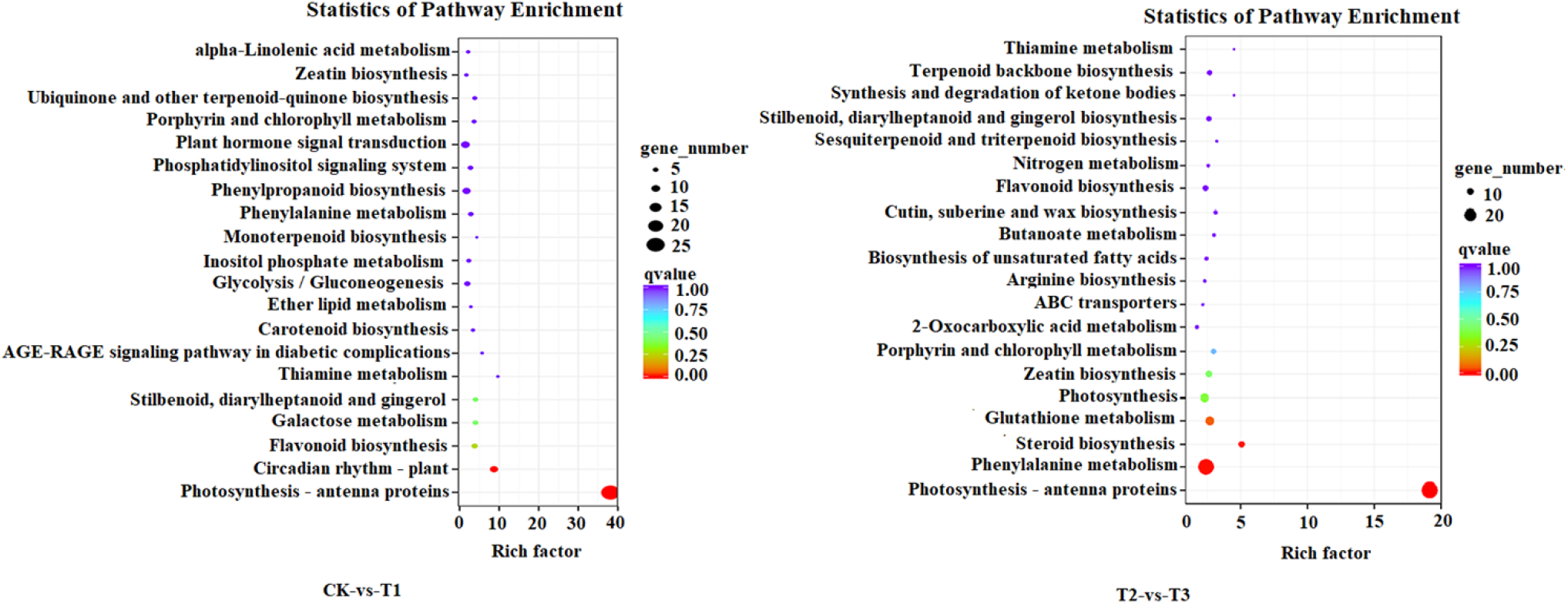
KEGG-enriched DEGs in salt treatment, c-GMP treatment, and the combination of both. Functional analysis of differential genes was also performed using GO analysis. Differentially expressed genes could be divided into three categories: biological processes, molecular functions, and cellular components. Higher GO terms were metabolic process, catalytic activity, cellular process, single-organism process, cell, cell part, and binding (Figure 4A, Table S6) in CK-vs-T1. Higher GO terms were metabolic process, binding, single-organism process, cell, cell part, catalytic activity, and cellular process (Figure 4B, Table S7) in T2-vs-T3.

**Figure 4.**
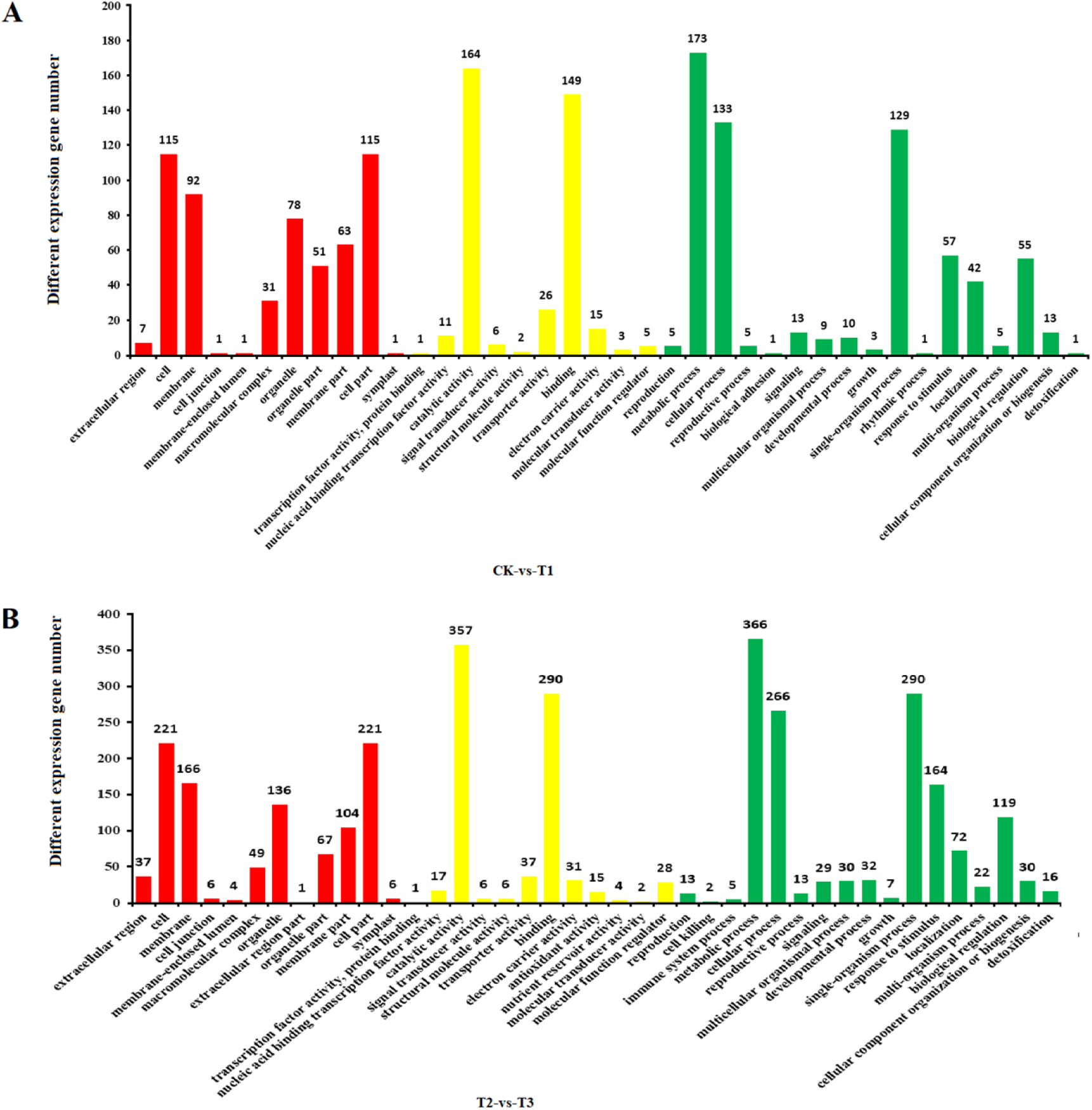
Visualization of go enrichment terms related to salt stress and c-GMP reaction. Functional classification of GO terms obtained using WEGO software and summarized using REVIGO. Terminology associated with DEGs with c-GMP treatment (A) and c-GMP treatment under salt stress (B)

### 2.5 Transcription factors analysis and SNP (single nucleotide polymorphism) analysis

Transcription factors (TFs) play an important regulatory role in the growth and development of plants and in response to changes in the external environment. TFs are the key link in regulating plant responses to drought, high salt, low temperature, hormones, and pathogens. The RNA-seq results showed that many TFs were differentially regulated in young tomato leaves under different treatments; we identified differential expression of 67 transcription factor families in tomato (Figure 5, Table S8). Some families were very sensitive to c-GMP-mediated salt stress, including AP2, b-HLH, C2H2, MYB, and NAC. The c-GMP-induced expression of these transcription factor gene family plays a key role in tomato resistance to salt stress.

**Figure 5.**
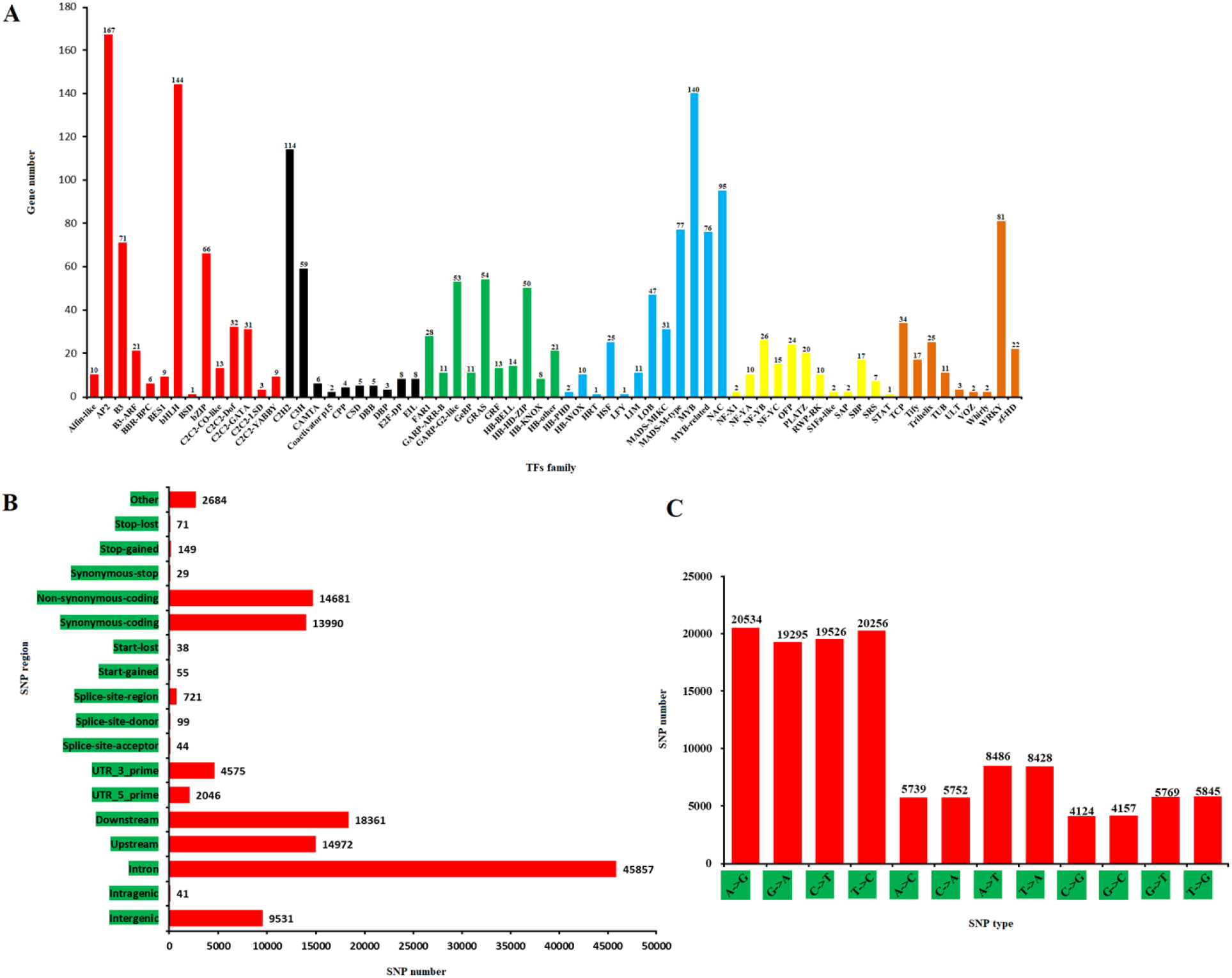
Summary of statistical analysis of transcription factors (TFs) and SNPs with different treatment. (A) 67 Differential expression TFs; (B) SNP distribution area; (C) SNP type

SNPs were identified in the assembled transcriptomes. We found 127,911 SNPs in the transcriptome, with 79,611 being transitions, and 48,300 transversions. The frequency of A/G (31.13%) and C/T (31.10%) were the highest in the transition type, and the highest frequency of A/T was 13.22% in the transversion. Most SNPs were located in non-coding regions; there were only 28,958 SNPs in the coding regions, and 45,857 SNPs in the exon regions (Figure 5B, 5C, Table S9). The characterization of SNP information in transcriptome data in this study is a feasible and effective method for the development of SNP molecular markers, and it also provided a feasible reference for the analysis of specific traits in the future.

#### 2.6.1 Exogenous c-GMP induced genes associated with carbohydrate metabolism

Compared with CK, T1 and T2, exogenous c-GMP induced carbohydrate-related metabolic DEGs under salt stress (Table S11, Table S12). Among them, the 2,3-diphosphoglycerate-dependent phosphoglycerate mutase encoding gene (ID: solyc04g072800.2) was up-regulated, and the glyceraldehyde 3-phosphate dehydrogenase (GAPDH, EC: 1.2.1.9) encoding gene (ID: solyc01g098950.1) was down-regulated in the T3 treatment, which promoted accumulation of glyceraldehyde 3-phosphate.

Non-phosphorylated GAPDH (ID: solyc07g005390.2) is a reaction enzyme that activates glycolysis “bypass”, with the assistance of NADP^+^. It can catalyze glyceraldehyde 3-phosphate to glycerate 3-phosphate; the NADPH in the process ensured a smooth flow of carbon source under salt stress. In addition, the gene (EC: 2.3.1.9, ID: solyc07g045350.2) encoding acetyl-CoA was up-regulated. Acetyl-CoA was protected by the TCA, and it was involved in amylose and sucrose metabolism, amino sugar and nucleotide sugar metabolism and mutual conversion of pentose and glucuronic acid. To demonstrate the effect of exogenous c-GMP on carbohydrates under salt stress, the content of amylase and soluble sugar was determined (Figure 6). Compared to T2, the starch content in the T3 treatment decreased by 10.37%, and the soluble sugar content increased by 32.51%. This suggests that exogenous c-GMP enhanced the osmotic adjustment capacity by promoting decomposition of starch, thus diminishing salt stress in tomato.

**Figure 6.**
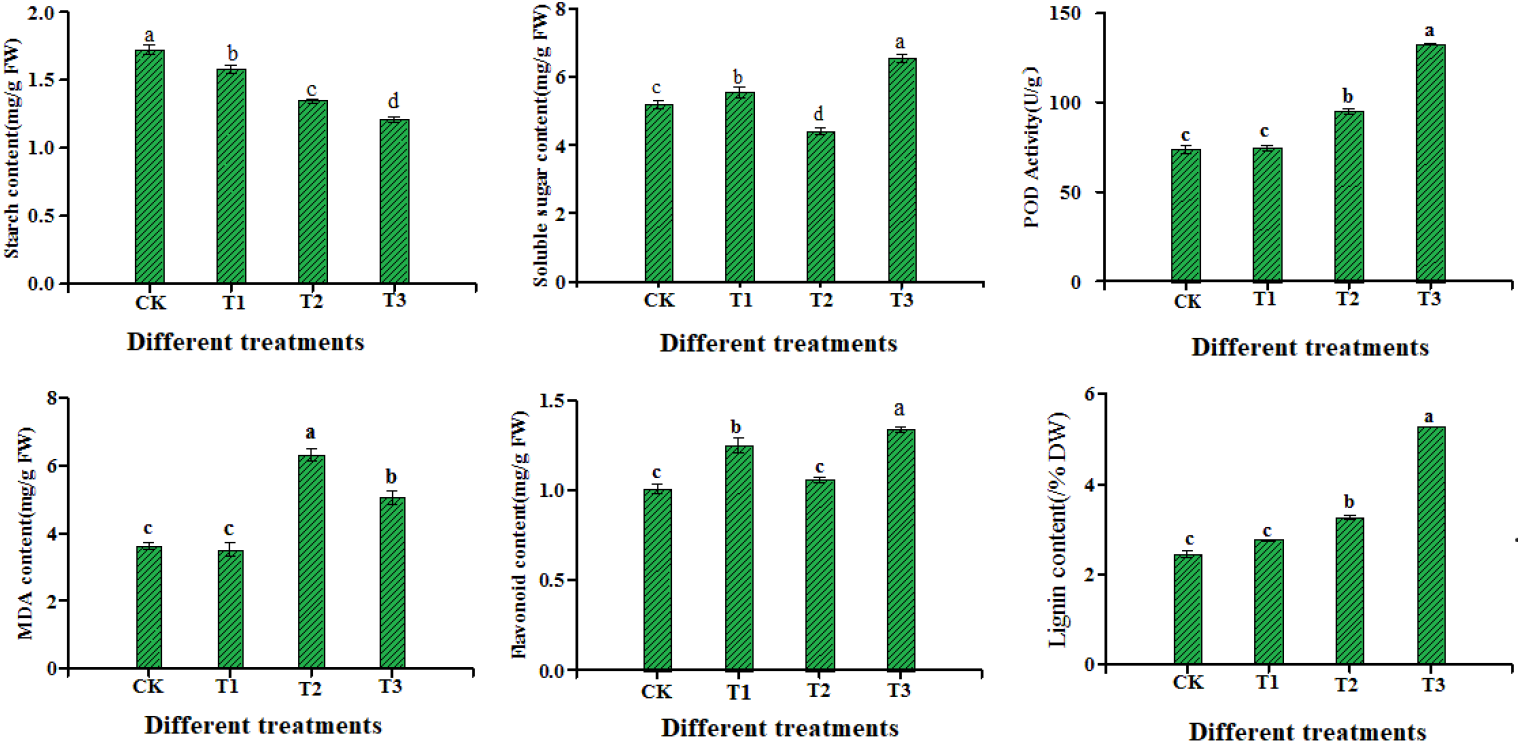
Physiological indexes of tomato seedlings in salt stress, c-GMP induction and GMP treatment under salt stress. The error bars represent the standard error of the mean. Significant differences between treatments at p ⩽ 0.05 are indicated by asterisks.

The genes coding for polygalactosidase (PG, EC: 3.2.1.67, ID: solyc07g041650.1) and pectin lyase (PL, EC: 4.2.2.2, ID: solyc05g014000.2) was down-regulated in T3, ensuring the protective effect of pectin in the cell wall, enhancing the salt tolerance of tomato. In addition, among pectinesterase DEGs, three were up-regulated (PMEs, EC: 3.1.1.11, ID: solyc03g123630.2, solyc09g075330.2 and solyc10g083290.1), and two (ID: solyc01g07918 0.2, solyc02g075620.2) were down-regulated. PMEs promoted a degrease in pectin and enhanced tomato cell buffering capacity to cope with ion changes caused by salt damage. The β-D-xylosidase 4 gene encoding cellulase (BXL4, EC: 3.2.1.37, ID: solyc10g047030.1) and the β-glucosidase gene (β -G, EC: 3.2.1.21, ID: solyc10g045240.1) were down-regulated in T3, slowing down the degradation rate of cellulose, enhancing the mechanical strength of the cell wall and improving the salt tolerance of tomato.

#### 2.6.2 Exogenous c-GMP induced genes associated with phenylpropanoid metabolism

Exogenous c-GMP regulated DEGs in phenylpropanoid-related metabolic pathways in plants exposed to salt stress. Phenylalanine metabolism, stilbenoid, diarylheptanoid and gingerol biosynthesis, and flavonoid biosynthesis were regulated in the T3 treatment (Table S11, Table S12). Some key genes involved in lignin and flavonoid synthesis, such as a gene encoding phenylalanine lyase (PAL, EC: 4.2.1.24, ID: solyc05g056170.2), were up-regulated in the T3. Four genes encoding caffeoyl-CoA-methylreductase (CCoAOMTs, EC: 2.1.1.104, ID: solyc02g093230.2, solyc02g093250.2, solyc02g093270.2, and solyc04g063210.2) were up-regulated in the T3 and down-regulated in the T2 treatment.

DEGs encoding peroxidases (PODs, EC: 1.11.1.7, ID: solyc01g067870.2, solyc02g080530.2, solyc10g076190.2, solyc10g076210.2, and solyc10g076220.2) were up-regulated in the T2 and down-regulated in the CK, T1 and T3. In addition, 58.33% of PODs (ID: solyc01g006300.2, solyc01g105070.2, solyc02g079500.2, solyc02g094180.2, solyc03g044100.2, solyc04g071890.2, and solyc09g007520.2) were up-regulated in the CK, T1 and T3 and down-regulated in the T2 treatment. POD is a key enzyme in the final step of lignin and flavonoid synthesis, and as a scavenger of ROS plays an important role in salt tolerance. We determined the POD activity and lignin and flavonoid content in each treatment (Figure 6). Compared with T2, the POD activity increased in T3 group by 28.38%, and the lignin and flavonoid contents increased by 38.07% and 20.89%, respectively. This suggests that cGMP could alleviate the oxidative damage caused by salt stress in tomato seedlings by regulating flavonoids and antioxidant enzymes, and promoting the biosynthesis of lignin, thus strengthening the hydrophobicity and mechanical strength of the cell walls to alleviate osmotic stress in salt-treated tomato.

#### 2.6.3 Exogenous c-GMP induced genes associated with fatty acid metabolism

Fatty acid desaturases (FADs, EC: 1.14.19.61.14.19.22) are key enzymes in the metabolism of fatty acids, regulating the composition and content of fatty acids in cell membranes (Tables S11 and S12). Compared with CK, T1 and T2, in the T3 treatment exogenous c-GMP induced up-regulation of four genes encoding FADs (ID: solyc04g040120.1, solyc12g049030.1, solyc12g100250.1, and solyc12g100260.1), maintaining membrane structure and composition, thus alleviating the inhibition of tomato growth by salt stress. During the fatty acid oxidation process, a gene encoding acetyl CoA (ID: solyc07g045350.2) was up-regulated in T3 treatment; acetyl CoA produced CO2 through TCA cycle, and promoted ATP synthesis to facilitate tomato growth and antioxidant system, effectively alleviating salt stress through indirect action.

Aldehyde decarboxylase (AD, EC: 4.1.99.5) is one of the key enzymes in the process of cuticle formation and wax biosynthesis. The gene encoding AD (ID: solyc12g100270.1 and solyc08g044260.2) was 50% up-regulated in T3 treatment, reducing dehydration to resist water losses caused by salt stress.

To demonstrate the stability of the membrane lipids, we determined the malondialdehyde (MDA) content (Figure 6). The MDA was significantly lower in the T3 than T2 treatment, suggesting that exogenous c-GMP effectively alleviated the membrane lipid peroxidation in tomato exposed to salt stress.

#### 2.6.4 Validation of Differential Genes by Quantitative RT-PCR Analysis

To verify the credibility of the RNA-seq results, relative expression levels of 9 DEGs involved in carbohydrate metabolism, phenylpropanoid metabolism, and fatty acid metabolic pathways were examined by real-time quantitative PCR (qRT-PCR) using independent samples (Figure 7). The test data indicated that 9 DEGs showed almost identical trends between qRT-PCR results and RNA-seq analysis, which demonstrated the reliability of RNA-seq sequencing data (Table S13).

**Figure 7.**
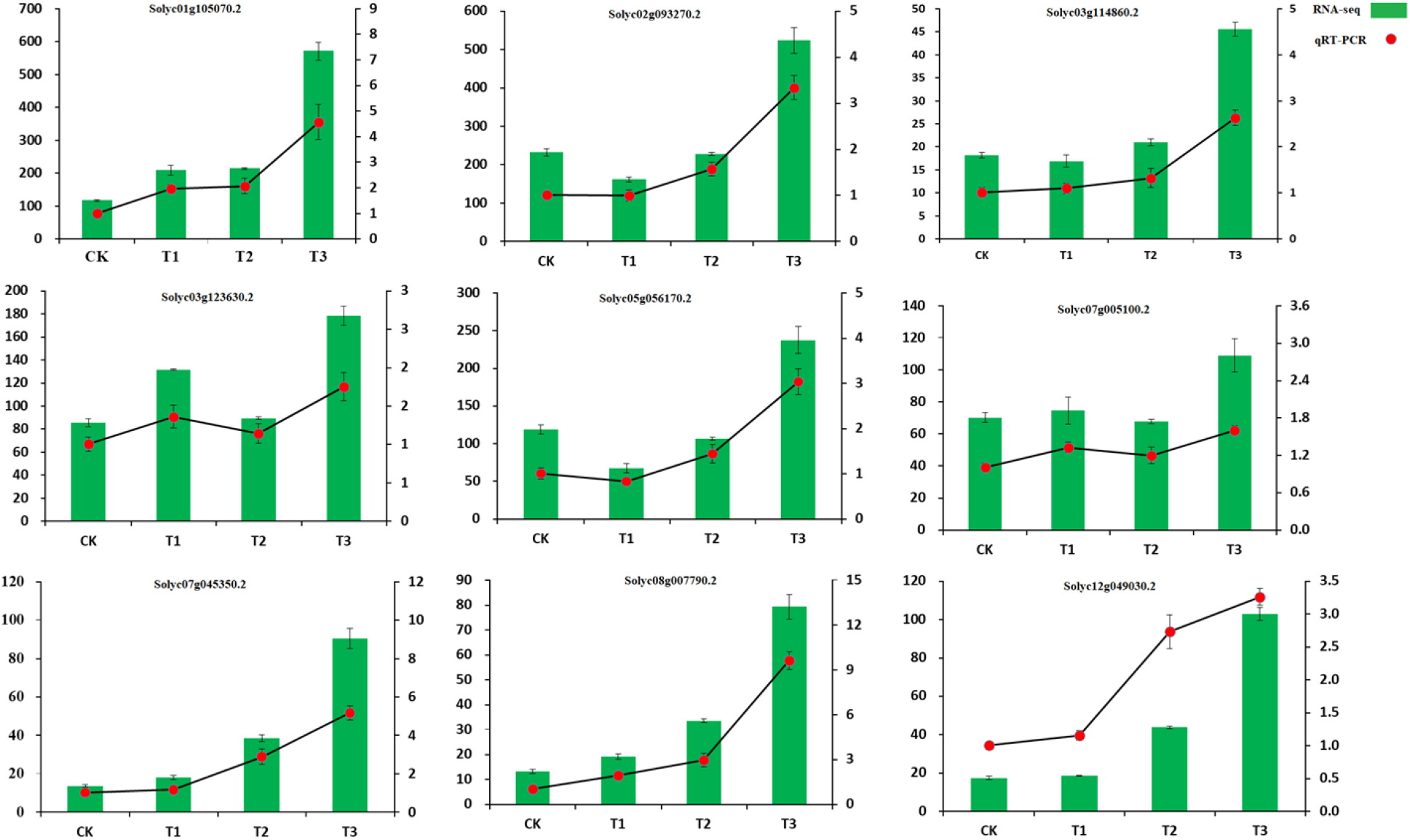
q-RT-PCR verification of c-GMP treatment of 9 differential genes under salt stress. Bars with standard errors represent relative expression levels determined by q-RT-PCR from three independent biological replicates using the 2^-ΔΔCT^ method (right y-axis). Broken lines indicate transcript abundance change (log2 fold) according to the RPKM value of RNA-Seq (left y-axis).

## Discussion

Salt stress is one of the most serious abiotic stresses affecting crop yield and quality. Most studies have researched the long-term response of plants to salt stress, focusing on the response of different organs, including leaves, roots and calli. For instance, root growth and seed germination of *Arabidopsis thaliana* were severely inhibited under salt stress [40]. Understanding the mechanisms of salt tolerance in tomatoes may allow effective use the saline-alkali soils to cultivate tomatoes, promote the green industry and develop strategies of growing vegetables.

c-GMP, as an important messenger substance in animals and plants, has been shown to promote plant development and enhance responses to abiotic stresses [9], playing a key regulatory role in *Arabidopsis* seed germination under salt stress[41]. To date, most studies have focused on the interaction of c-GMP with other signaling molecules, such as NO, to enhance plant tolerance. Yet, there is little information on the role of exogenous c-GMP in promoting tomato salt resistance. Recently, researchers found that salt stress activates the c-GMP signaling pathway [42], and that c-GMP regulates the transport of monovalent cations (Na^+^ and K^+^) in *Arabidopsis* [43]. However, the regulatory mechanisms of c-GMP in plant resistance to salt stress and related signal transduction processes are still unclear. Understanding the molecular mechanisms of exogenous c-GMP in abiotic stress tolerance is important for breeding programs aimed at producing salt-tolerant varieties. Therefore, we conducted transcriptome analysis of tomato in response to salt stress and c-GMP treatment. The study detected 1884 stress-responsive DEGs in the salt treatment. We further found 537 and 1043 DEGs in the comparison CK-vs-T1 and T2-vs-T3. These results indicated the diversity and complexity of exogenous c-GMP-regulated genes in tomato seedlings under salt stress.

Studies have shown that the AP2 transcription factors are widely present in plant genomes, regulating plant growth and development, and playing a key regulatory role in plant response to salt stress [44]. In the present study, 182 AP2 transcription factors were differentially expressed. bHLHs can regulate plant responses to abiotic stress, and its role is very significant in plant stress resistance. Protein of b-HLH family (*bHLH30*) was up-regulated in the treatments with salt alone as well salt + c-GMP, with the former showing greater expression. In particular, the WRKY gene family is regarded to be actively involved in salt stress responses, and complex regulatory mechanisms of WRKY have been observed under different stress conditions [45]. For example, under salt stress, *ZmWRKY33* in maize [46] and *GhWRKY39* in cotton [47] were up-regulated. In the present study, eight WRKYs genes were up-regulated under salt stress alone, but their expression levels returned to normal under salt stress + c-GMP. These results revealed that c-GMP treatment can inhibit the expression of b-HLH and WRKY transcription factors under salt stress.

SNP molecular markers are highly specific genetic markers. With the wide application of high-throughput sequencing technology and a gradual decrease in the cost of detecting SNP markers, mining SNP sites based on transcriptome sequencing will make SNP molecular marker technology increasingly efficient. For many species, a large number of SNPs can be identified using next-generation high-throughput transcriptome sequencing technologies [48]. In recent years, SNPs have also been used successfully in research on plant genotyping, variety identification, and new genotype selection. Li [49] used 14 SNP markers to genotype 30 citrus varieties, distinguishing between varieties kinship. Distefano[50] identified 21 SNP markers for citrus and analyzed genetic diversity of 18 citrus varieties. However, few SNP molecular markers have been studied in tomato so far. Our study enriched the SNP information in tomato genome by mining SNP sites through transcriptome, and found the frequency of transitions was significantly (about 2-fold) higher than of transversions, which was similar to the ratio of SNP variation types in other plants and transcriptomes[51]. The statistic of SNP loci provides a reference for further research of SNP molecular marker development.

The photosynthetic rate in fresh leaves of plants was extremely weak when plants were under stress, with leaves reducing the water loss by closing the stomata, thereby blocking CO2 supply and reducing photosynthetic capacity [52–53]. In the present study, KEGG enrichment analysis was performed on all DEGs (CK-vs-T1 and T2-vs-T3). Under control conditions and salt stress, we found exogenous c-GMP induced, respectively, 26 and 28 DEGs enriched in the photosynthesis-antenna protein pathway, but were all down-regulated, suggesting that the amount of carbohydrates produced by photosynthetic carbon fixation would not meet the metabolic requirements.

In the T3 treatment, exogenous c-GMP governed the down-regulation of the gene encoding sucrose synthase in the starch and sucrose metabolism pathway, whereas the gene encoding β-fructofuranosidase (Table S10, Table S12) was up-regulated; The reduction in carbon allocated to sucrose synthesis led to a decrease in starch content, which promoted the accumulation of soluble sugars, such as fructose and glucose, to buffer the damage caused by salt stress in tomato. The above results were consistent with study of arbuscular mycorrhiza increasing the accumulation of soluble sugars in maize roots and enhancing the tolerance to salt stress [75] The redox state of plants can regulate multiple functions of cells; it is a prerequisite for plants to resist adverse environmental conditions and protect themselves from oxidative stress. The key pathway of plant carbon cycle includes glycolysis and TCA cycle. GAPDH plays a central role in the cell carbon cycle and is one of the key enzymes for maintaining energy supply for metabolic processes. In the present study, the genes encoding GAPDHs were down-regulated, whereas NADPH accumulation guaranteed the access to carbon source under salt stress, which might also reduced the toxicity of free ammonia produced by the hydrolysis of proteins during ionic stress in tomato. The stored nitrogen and carbon sources would provide energy for restoring growth after stress was removed.

Secondary metabolites play an important role in plant stress tolerance [54]. Transcriptional analysis of various genes in the phenylpropanoid pathway under salt stress was carried out in plants such as *Arabidopsis thaliana* [32], poplar [55], and black pepper [56]. These analyses indicate that this pathway is incorporated into the defense response [57]. In the present study, genes were identified in the major branch of the phenylpropane pathway, which is the biosynthesis of lignin and flavonoids. The cell wall is an active structure that mediates the critical stages of plant development, especially in response to stress signals, which can induce differential expression of the coding genes of pectin and cellulose to rapidly adapt to adversity [58–59]. Exogenous c-GMP regulated DEGs associated with carbohydrates in the phenylpropane metabolic pathway under salt stress compared to normal conditions. Among them, 3 of the 5 *PMEs* in the T3 treatment were up-regulated, and the accumulation of pectin provided an important buffering capacity for tomato cells to cope with the ion changes caused by salt damage. Down-regulation of the gene encoding cellulase slowed the degradation rate of cellulose, and this result was consistent with the analysis of the cloning and expression patterns of the *FmMUR5* gene of ash [60]. In addition, PAL and most of CCoAOMT and PODs (Table S10, Table S12) were up-regulated to promote the synthesis of lignin and flavonoid. This indicated that exogenous c-GMP enhanced stress tolerance by regulating the constituents of tomato cell walls to enhance mechanical tolerance and hydrophobicity, and transmitted cell stress signals.

As low molecular weight secondary metabolites, flavonoids are antioxidants that respond effectively to salt stress. Plants evolved macromolecules to act as osmotic regulators under stress and play an important role in maintaining the osmotic balance of cell fluids [61].

Previous results showed that ROS (reactive oxygen species) was signal molecules involved in a cucumber response to low temperature and paraquat stress [62]. Excessive accumulation of ROS in cell compartments is a common phenomenon caused by salt stress in plants; plants have developed antioxidant enzymes such as superoxide dismutase (SOD), POD, GPX, and catalase to detoxify free radicals. MDA is a product of lipid peroxidation, and the extent of membrane damage can be reflected in the concentration of MDA in the stressed tissue [63]. POD is involved in removing excess ROS, which helps plants withstand stress conditions [64]. POD is not only an important substance in the final step of lignin synthesis, but also acts as an antioxidant enzyme to scavenge ROS. In the present study the content of soluble sugar, and activity of protective enzyme (POD and MDA) were higher in T3 than T2 treatment. These results indicated that exogenously administered c-GMP protected plants from oxidative stress by enzymatic and non-enzymatic ROS scavengers under salt stress.

Salt stress affects plant growth mainly by reducing the water absorption capacity of plants. In order to satisfy their own metabolic needs, plants usually close the stomata to reduce water loss, or accumulate proline to reduce permeability and water loss, and also accumulate wax on the surface of leaves. In addition, the cuticle reduces non-porous water loss [65]. The fatty acid metabolism in plants can also ensure the good fluidity of the membranes by desaturation, mobilization and regulation of key enzyme activities, providing an excellent environment for protein activity. Thus, dealing with adverse damage, the realization of this process depends on the effective regulation of FADs. Genes encoding FADs have been described briefly in rice [66] and maize [67], and the regulation of their expression by abiotic stresses occurs at the transcriptional and post-transcriptional levels[68–69]. In the present study, compared with normal conditions, exogenous c-GMP induced significant accumulation of DEGs in tomato in the fatty acid metabolism pathway under salt stress. Further analysis revealed that c-GMP-regulated wax synthesis was based mainly on prolonged fatty acids in vitro and the decarbonylation pathway in the process of derivatives, which was studied in pea (*Pisum sativum*) and chlorella (*Botryococcus braunii*) [70–71]. In addition, studies in *Arabidopsis* found that decarbonylation pathways were enhanced. The waxy component accounted for more than 90% of the total waxy substance, and the decarbonylation pathway repression caused a significant reduction in the total wax content of the *Arabidopsis* stem [72]. In the T3 treatment, exogenous c-GMP induced the AD gene that was up-regulated under salt stress, and various components of the plant epidermis wax were hydrophobic, indicating that c-GMP was likely to reduce non-stomatal water loss by regulating the synthesis of wax in fatty acid metabolism.

Based on our findings, a network of key genes induced by c-GMP under salt stress were involved with metabolism of carbohydrate, fatty acid and phenylpropanoid as mapped (Figure 8). The gene expression values were based on the KEGG pathway approach. This could provide many insights into molecular mechanisms of c-GMP induced in adaptation of tomato seedlings to salt stress.

**Figure 8.**
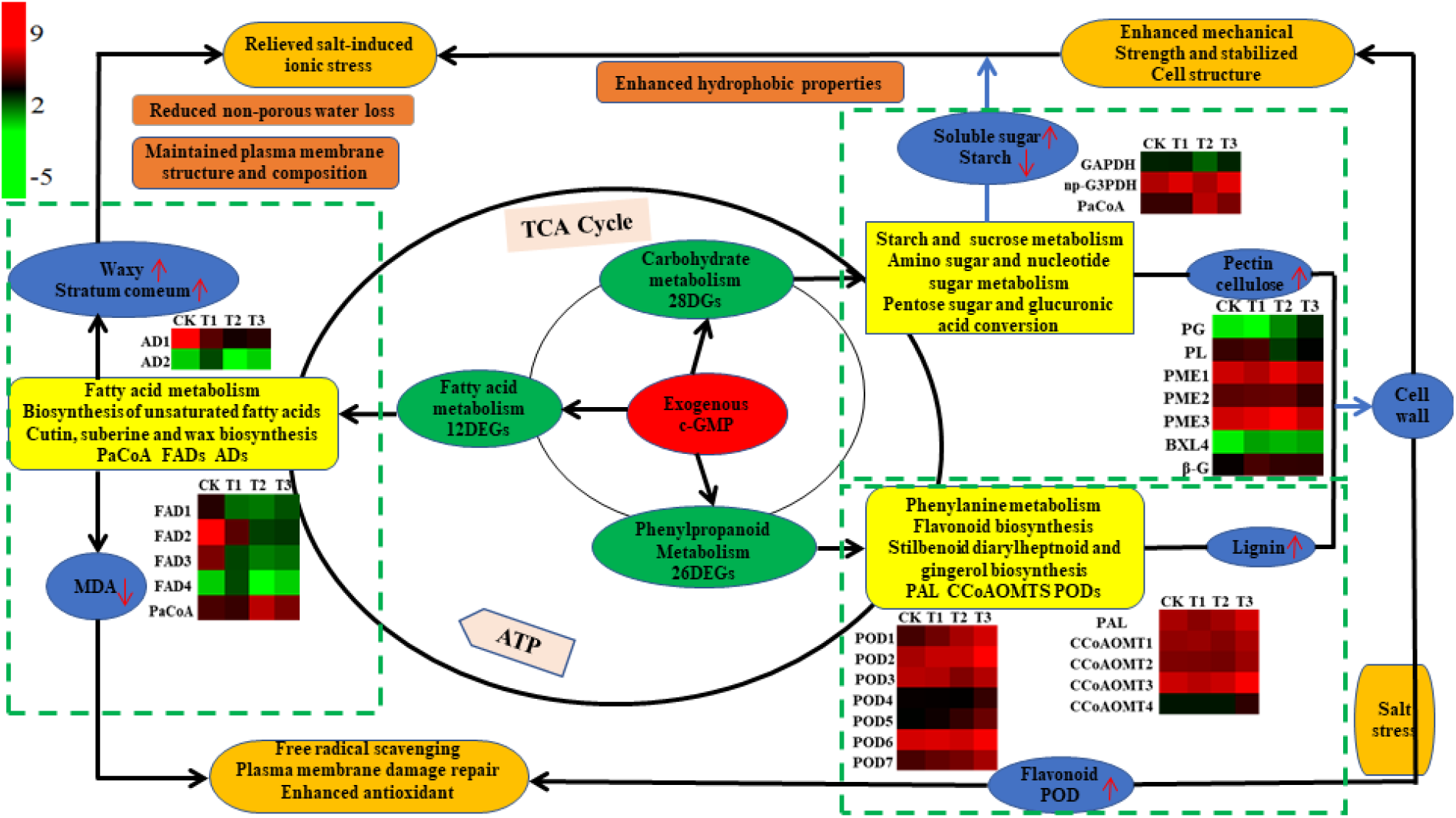
The network of key genes induced by c-GMP under salt stress in the metabolism of carbohydrate, fatty acid and phenylpropanoid.

## Conclusions

Tomato seedlings can overcome salt stress by regulating carbohydrate, phenylpropanoid and fatty acid metabolism through inducing differential expression of related genes, in addition to altering the composition of the cell wall to enhance hydrophobic properties (Figure 8). Exogenous application of c-GMP can improve the osmotic adjustment, enhance the antioxidant defense, decrease membrane lipid oxidation, and reduce the non-stomatal water loss to strengthen the salt tolerance of tomato seedlings. This study provides a solid foundation for better characterization of salt-resistant genes, and also gives insights into salt tolerance of tomato seedlings, and may contribute to the field production of tomato in the arid and semi-arid regions of the world.

